# Neural processing of linguistic speech features and their modulation by selective attention in users of hearing aids and cochlear implants

**DOI:** 10.64898/2026.11.12.765375

**Authors:** Constantin Jehn, Alina Schüller, Niki Katerina Vavatzanidis, Anja Hahne, Tobias Reichenbach

**Affiliations:** Department Artificial Intelligence in Biomedical Engineering (AIBE), Friedrich-Alexander-Universität Erlangen-Nürnberg (FAU), Nürnberger Strasse 74, 91052 Erlangen, Germany; Ear Research Center Dresden (ERCD), University Hospital Carl Gustav, Carus, Technische Universität Dresden, Dresden, Germany

## Abstract

Understanding speech relies on the neural processing of acoustic and linguistic features, which can both be analyzed using Temporal Response Functions (TRFs). While both types of features have been the focus of numerous studies in typical-hearing individuals, much less is known about the neural processing of continuous speech for people with hearing loss, particularly how hearing-impaired listeners process word-level features in noisy environments, where they typically struggle. We recorded EEG from hearing aid (HA, n=29) and cochlear implant (CI, n=24) users, and an age-matched typical-hearing (TH) control group (n=29), while participants listened to audiobook segments in quiet and with a competing speaker, using their personal hearing devices (CI and HA). A hierarchical encoding framework incrementally added word onset, word frequency, and word surprisal as predictors of the EEG signal. Attentional modulation was assessed by comparing encoding scores for attended versus ignored speech. All three groups showed clear neural responses to word onset, frequency, and surprisal for speech in quiet. Surprisal tracking emerged at 400 ms in TH participants, 500 ms in HA users, and 600 ms in CI users, weakest in the CI group. All three groups exhibited attentional modulation of surprisal tracking. Word frequency yielded significant encoding gains in quiet only for TH and HI listeners, not CI users. We show for the first time that CI users exhibit significant, attention-modulated neural responses to word surprisal. More broadly, surprisal tracking was present across all hearing-impaired groups but weakest in CI users, the cohort with the greatest difficulty understanding speech in noise.

## 1 Introduction

Hearing loss contributed to 43–54 million years lived with disability globally in 2019, making it the third largest cause of disability [1]. A main difficulty for affected individuals is to follow and participate in conversations in noisy environments [2, 3], which may contribute to the association found between hearing loss and decreased social and mental well-being [4, 5].

Hearing aids treat moderate to severe hearing loss by amplifying incoming sound to improve audibility [6]. However, users struggle in noisy environments, showing increased listening effort and reduced speech comprehension [7]. Cochlear implants (CIs) treat severe to profound hearing loss by stimulating the auditory nerve, bypassing the auditory periphery [8]. CIs restore speech understanding in quiet, but speech comprehension in noise poses a bigger challenge than for hearing aid users: CI processing preserves temporal fine structure poorly, and CI users show reduced spatial release from masking [9, 10, 11, 12, 13].

Understanding speech in noise depends on central auditory and cognitive processes [14]. Attention filters a target speech stream from other acoustic signals [15, 16]. This selective processing appears in neural speech tracking, i.e., cortical tracking of speech features such as the speech envelope [17, 18]. Encoding models quantify this tracking by characterizing how speech features predict neural responses [19], while decoding models reconstruct speech features from neural data. Both approaches show larger tracking of a speech stream when attended than when ignored [18].

Multiple studies examined selective envelope tracking in hearing-impaired versus typical-hearing individuals. Fuglsang et al. [20] and Decruy et al. [21] reported enhanced neural tracking of attended speech in hearing aid users, suggesting compensatory cortical gain under complex listening. In both studies, participants wore no clinically fitted devices. Instead, they listened to the original sound with frequency-specific amplification. Petersen et al. [22], in contrast, found a smaller attended-ignored difference in hearing aid users, with stimuli streamed directly to research devices rather than presented in a free field. In a free-field study using clinically fitted devices, we found cortical envelope tracking and its attentional modulation in hearing aid users comparable to typical-hearing individuals, despite increased listening effort [23]. This suggested that higher listening effort manifests in neural responses beyond cortical envelope tracking.

In CI users, the selectivity of the neural response to the speech envelope has been shown to be significantly reduced compared to both typical-hearing participants and hearing aid users [24, 25]. We found that the attentional modulation of the cortical speech tracking in CI users was also much smaller than in the other two groups [23].

The speech envelope is an acoustic feature of speech, and its cortical tracking reflects low-level acoustic processing. Speech and linguistic features, such as phoneme identity, word onsets, and word surprisal, can likewise be tracked in neural activity and linked to higher-level linguistic processing [26, 27, 28, 29]. Word surprisal, i.e., unexpectedness given prior context, can be quantified and its neural tracking peaks around 400 ms post-onset, consistent with the N400 ERP component [27, 30]. In competing speech paradigms, typical-hearing individuals show suppressed surprisal tracking for ignored speakers, indicating that surprisal depends on active speech comprehension [31, 32, 27, 30].

Only limited work has examined how hearing impairment affects linguistic-level speech tracking. One study on word-level segmentation, treating hearing ability as a covariate for cognitive decline, found that neural encoding accuracy decreased with increasing hearing loss [33]. As a single-speaker paradigm was used, competing-speaker effects were left open. Another study fitted hearing aid users with a standardized study device and tested phoneme encoding in a competing-speaker paradigm [34]. Attentional modulation was preserved, but phoneme-onset encoding was similar for target and ignored speakers, suggesting a failure to suppress irrelevant speech at this acoustic-phonetic stage. Without a typical-hearing control group, it remains unclear whether this reflects hearing loss per se or cocktail-party listening more generally.

No study has yet assessed neural tracking of linguistic speech features in CI users. For hearing aid users, data remain scarce and lack systematic comparison with typical-hearing individuals for competing speech.

The present study aims to close this gap. For the first time, it investigates linguistic speech tracking in hearing aid and CI users alongside typical-hearing controls, in both single-speaker and competing-speaker conditions. Such an assessment of higher-level linguistic speech processing may help explain behavioral findings, such as increased listening effort in hearing aid users, that neural tracking of lower-level acoustic features may fail to reveal.

## 2 Methods

### 2.1 Participants

The current study is a re-analysis of previously published data [23] on auditory attention decoding in CI users, HA users, and typical-hearing individuals. In total, 95 participants were recruited for the original study. All participants provided written informed consent, and the experiment was approved by the science ethics committee of Technische Universität Dresden (protocol SR+BO-EK-47022024) on May 21, 2024. All procedures were conducted in accordance with the Declaration of Helsinki.

Both the CI and HA participants used their own everyday device settings during the experiment, allowing cortical tracking to be assessed under ecologically valid conditions.

#### 2.1.1 Cochlear implant group (CI)

The CI group comprised 30 bilateral CI users, of whom 24 were included in the analysis after excluding six participants due to technical recording issues (n=4) or inability to complete the study (n=2). CI experience ranged from 2 to 28 years (median = 10 years). The majority of CI users had post-lingual hearing loss (n=20), with four participants classified as pre- or perilingual. To ensure sufficient task performance in the selective attention paradigm, participants were required to score at least 60% on the Freiburg monosyllabic word recognition test in quiet at 65 dB for their better-performing ear [35]. Speech comprehension in noise was additionally assessed at the sentence level using the HSM sentence test (65 dB signal, 60 dB noise, free field) [36].

#### 2.1.2 Hearing aid group (HA)

The HA group comprised 29 bilateral HA users, who completed the Freiburg monosyllabic word test in quiet (65 dB) and the HSM sentence test (65 dB signal, 60 dB noise) with hearing aids on, in a free-field environment. Word test data from five participants administered via headphones were excluded from analysis due to non-compliance with standard HA protocols [37]. All remaining participants completed the monaural word test with the contralateral ear occluded. Unaided pure-tone audiometry scores were obtained from hearing-care professionals or measured before the study, within two years for 27 participants (median: 8 months, range: 0–21 months; two outliers at 64 and 85 months).

#### 2.1.3 Typical-hearing group (TH)

An age-matched control group of 33 typical-hearing individuals was recruited, with inclusion requiring audiometric thresholds better than 25 dB per ear verified by pure-tone audiometry (PTA4). Four participants were excluded for not meeting this criterion, leaving a final TH group of 29 individuals.

#### 2.1.4 Distributions of sex, age, and hearing performance across groups

The three groups were matched in age (Kruskal–Wallis: *H*(2) = 1.03, *p* = 0.597, *η*^2^*_H_* = 0.013) and sex distribution (*χ*^2^(2) = 0.100, *p* = 0.951), as summarized in Table 1. All TH participants met the audiometric inclusion criterion (PTA4 *<* 25 dB; median = 8.75 dB), while the HA group showed - as expected - substantially elevated unaided thresholds (median = 46.9 dB; MWU: *p <* 0.001, *r*_rb_ = 1.0). Speech comprehension was assessed in quiet with the Freiburg monosyllabic word test and in noise with the HSM sentence test (65 dB signal and 60 dB noise). HA and CI users performed comparably in quiet (both medians = 85%; MWU: *p* = 0.796, *r*_rb_ = 0.05). Group differences emerged specifically in noise: HA users scored markedly higher than CI users on the HSM sentence test (medians: 65.1% vs. 23.0%; MWU: *p <* 0.001, *r*_rb_ = 0.64), providing the first indication that perceptual differences between device groups manifest under adverse listening conditions. Speech comprehension tests were not administered to TH participants, as pilot testing confirmed ceiling performance for both tests.

**Table 1:**
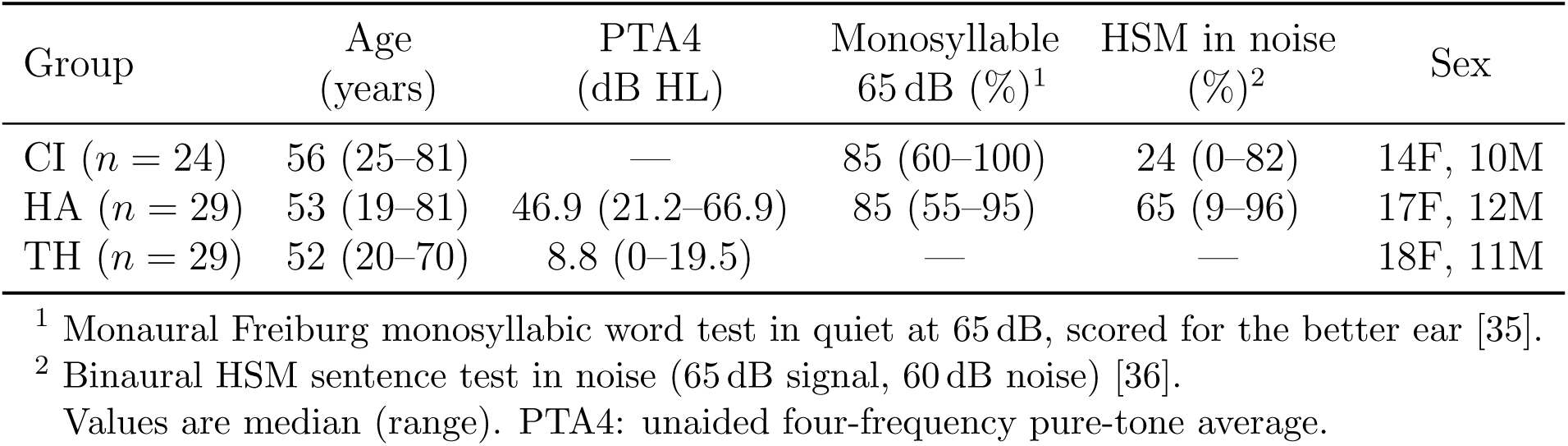
Demographic and audiological characteristics by group.

### 2.2 Stimuli and experimental procedures

Two German audiobooks served as stimuli: a male-narrated story (J.R.R. Tolkien, narrated by Gert Heidenreich) and a female-narrated story (Christine Ritter, narrated by Vera Teltz) [38, 39]. The male narrator’s speech rate was adjusted to 90% to match the female narrator. The two voices were acoustically distinct, with the female narrator showing higher fundamental frequency (164.5 *±* 46.3 Hz vs. 110.9 *±* 32.5 Hz), higher spectral centroid (4.27 *±* 2.87 kHz vs. 3.46 *±* 1.90 kHz), and shorter vocal tract length (14.2 cm vs. 16.7 cm), consistent with typical male–female voice differences. Both excerpts were loudness-normalised to *−*24 LUFS [40].

Stimuli were presented in a free-field environment via two loudspeakers positioned at 30*^◦^* of the left and right of the participant (Fig. 1 A). CI and HA listeners used their everyday device settings throughout. Each participant completed 20 trials of approximately two minutes each: eight single-speaker trials and twelve competing-speaker trials in which both audiobooks were presented simultaneously from spatially different loudspeakers. In the competing-speaker trials, a visual cue indicated the target speaker, who began 10 s before the distractor. Speaker identity and side of presentation were counterbalanced across trials.

**Figure 1:**
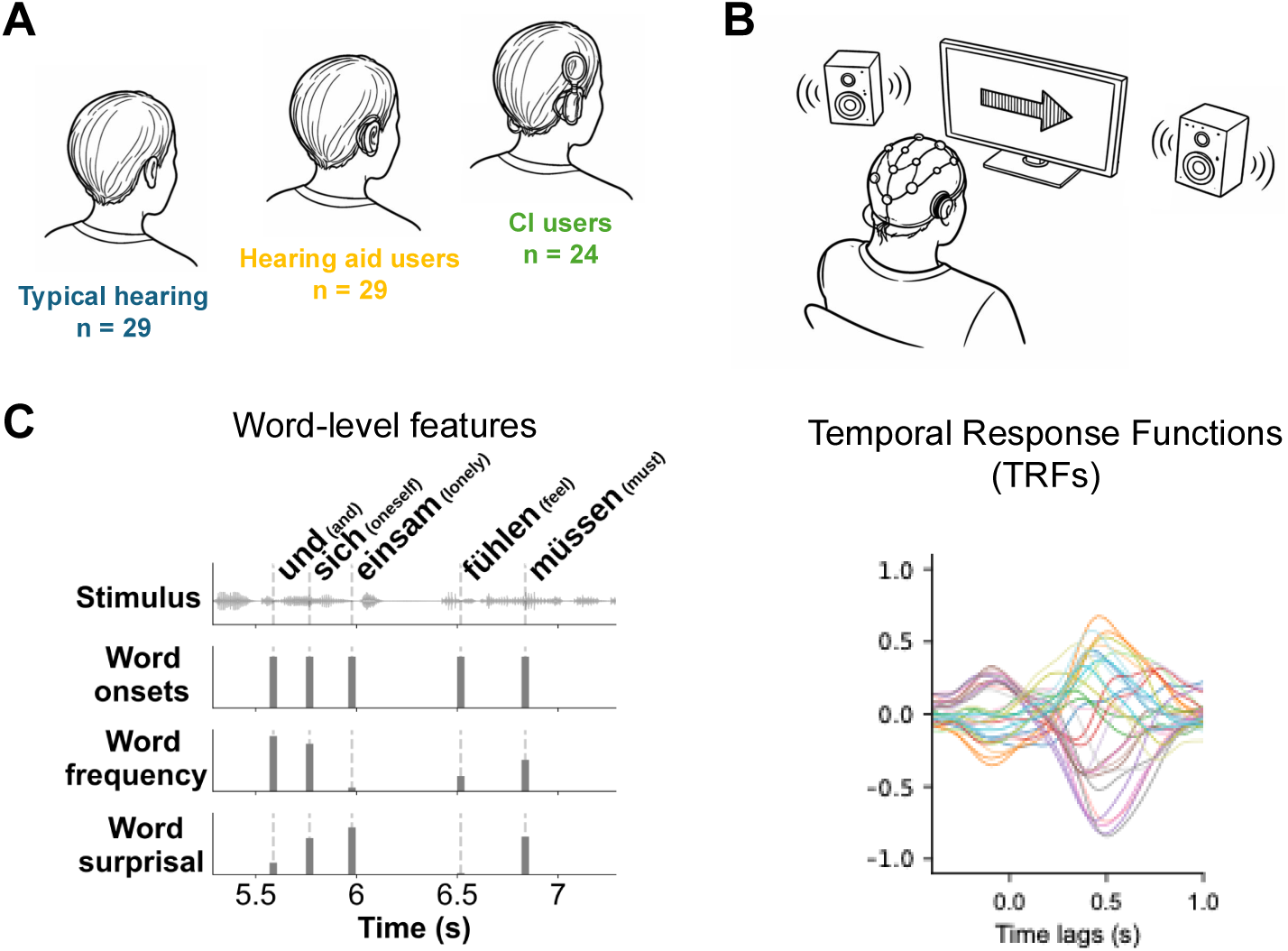
Overview of the experimental design and analysis pipeline. (**A**) Data were collected from 29 typical-hearing individuals, 29 hearing aid users, and 24 CI users. All three groups had a similar age distribution. (**B**) After an initial single-speaker paradigm, participants performed a selective auditory attention task while their brain activity was recorded with EEG. Participants heard two stories presented in a competing manner through two different loudspeakers, located to the left and the right. Hearing aid and CI users completed the task with their clinically fitted devices. (**C**) To study neural tracking of word-level features, we extracted word onsets, word frequency, and word surprisal. These were used to predict the EEG through Temporal Response Functions (TRFs). A hierarchical modeling approach then quantified the incremental predictive contribution of each word-level feature.

### 2.3 Behavioral measures

After each trial, participants answered two three-alternative forced-choice comprehension questions about the attended audiobook. The comprehension score was defined as the proportion of correctly answered questions. Subjective listening effort was rated once per condition and attended speaker using the ACALES scale (1 = no effort, 13 = extreme effort) [41]. The behavioral results are reported in full in [23]; a summary is provided in Table 2 in the Appendix.

### 2.4 EEG recordings

EEG was recorded using a 32-electrode actiCHamp system (BrainProducts GmbH, Germany) at 1 kHz with an online low-pass filter at 280 Hz. One electrode was placed below the eye for ocular artifact detection, leaving 31 channels for neural activity. Electrode impedances were kept below 20 kΩ throughout. In CI users, electrodes positioned between the implant and the coil were removed where necessary (mean: 1.7 electrodes per participant).

To synchronise the EEG with the audio stimuli, the presented audio signal was simultaneously recorded as two auxiliary channels via StimTrak adapters (BrainProducts GmbH, Germany). Temporal alignment was established offline by selecting the delay by maximising the Pearson correlation between the recorded and clean stimulus waveforms. The alignment was further validated against onset triggers.

### 2.5 Data preprocessing

#### 2.5.1 EEG data preprocessing

EEG data were preprocessed trial-wise using MNE-Python [42]. For all cohorts, a high-pass filter (0.3 Hz, 4th-order Butterworth IIR) was first applied to remove slow drifts. Independent Component Analysis (ICA) was then performed on each trial (15 components, random state fixed for reproducibility). Components were classified automatically using ICLabel [43], and components labeled as eye blink, eye movement, muscle artifact, cardiac activity, line noise, or channel noise were excluded.

As CIs introduce significant artifacts in EEG recordings [44, 45], a dedicated preprocessing step was applied to suppress their magnitude. Following [46], an ICA was computed using the InfoMax algorithm [47], separating up to 30 components. Each component’s similarity to the audio stimulus was quantified as the SNR of the cross-correlation peak around zero lag (-5 to 12 ms), expressed in dB. Components exceeding an SNR of 25 dB were removed, corresponding to 3% of all independent components. The effectiveness of this approach has been demonstrated independently [48].

For all cohorts, a common average reference was applied, followed by a notch filter at 50 Hz to suppress line noise, and a low-pass filter at 62 Hz (4th-order Butterworth IIR). Data were subsequently low-pass filtered at 4.5 Hz (4th-order Butterworth IIR) and downsampled from 1,000 Hz to 100 Hz. The low-pass cutoff was chosen to isolate the delta frequency band (1–4 Hz), in which cortical tracking of word-level linguistic features, including word onsets, word frequency, and word surprisal, has been predominantly observed [27, 31, 49]. Finally, each EEG channel was z-score normalized across time.

#### 2.5.2 Linguistic feature extraction

To investigate EEG responses to word-level features of continuous speech, three word-level predictors were extracted for each stimulus: word onset, word frequency, and word surprisal. All features were derived from the acoustic stimuli, which were presented in the single-speaker and the competing speaker paradigms. This resulted in word features for the single male speaker, the single female speaker, the competing male speaker when he was attended, as well as when he was the distractor, and lastly, the competing female speaker when she was attended, as well as when she was the distractor.

##### Word onsets

Before onset extraction, each audio file was time-aligned with its corresponding transcript using WebMAUS Basic [50, 51], an automatic forced alignment service provided by the Bavarian Archive for Speech Signals (BAS) at Ludwig Maximilian University of Munich. WebMAUS Basic takes an audio recording and a word-level transcript as input and returns a Praat TextGrid file containing time-aligned word and phoneme boundaries, derived via hidden Markov model-based acoustic modeling. The German language model was used for all alignments. The resulting TextGrid files served as the basis for all subsequent onset extraction.

Word onset and offset times were then extracted from these TextGrid files. The first interval tier was used, and all labeled intervals (i.e., non-empty marks) were treated as words. Onset and offset times were converted from seconds to samples using the sampling rate of 48,000 Hz, yielding a precise sample-level time series of word boundaries for each sentence.

##### Word frequency

Word frequency was obtained from SUBTLEX-DE [52], a large-scale German subtitle corpus providing frequency norms for German words. The database was queried case-insensitively (e.g., aggregating counts for “Gehen” and “gehen”) to yield a single frequency estimate per word form. For each word in the stimulus transcripts, the Zipf word frequency value (ZipfSUBTLEX; [53]) was extracted, which represents log-transformed word frequency on a scale from 1 to 7 and provides a more perceptually meaningful measure of lexical availability. Words absent from the database received a frequency of zero.

##### Word surprisal

Word surprisal was computed as a measure of contextual predictability using the pretrained German causal language model german-gpt2 [54], which is based on the GPT-2 architecture [55]. For each sentence, the transcript was tokenized using the model’s byte-pair encoding (BPE) tokenizer. Surprisal was calculated incrementally: for each token *t_i_* at position *i*, the model’s probability *P* (*t_i_|t*_1_*, …, t_i−_*_1_) over the vocabulary was obtained given all preceding tokens of that sentence as context. The negative logarithmic probability of the observed token was computed as:

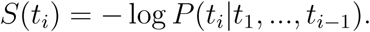

Because BPE tokenization may split a single orthographic word into multiple subword tokens (e.g. “Kinder” might be split into “Kind” and “er”), surprisal values were summed across all subword tokens belonging to the same word, yielding one surprisal estimate per word. This follows from the definition of surprisal as the negative log probability: the joint word probability is the product of token probabilities, and the negative logarithm converts this product into a sum.

Each word-level feature was represented as a sparse continuous time series at the EEG sampling rate of 100 Hz. For word onsets, a binary impulse train was constructed by placing a unit impulse at the sample corresponding to each word onset. For word frequency and surprisal, weighted impulse trains were constructed analogously, with each impulse scaled by the respective scalar value of that word. This encoding scheme was inspired by previous studies [27] and allows each feature to be modeled as a discrete event time series. Figure 1C (left) depicts a 2 s window of the constructed stimuli. Prior to model fitting, all three feature time series were independently z-score normalized across time.

### 2.6 Temporal Response Functions

To quantify the neural encoding of the three word-level features, we estimated Temporal Response Functions (TRFs) using a linear forward model. The TRF describes how the brain responds to each stimulus feature across a range of latencies, and can be understood as a linear filter that, when convolved with the feature time series, predicts the measured EEG activity. Formally, the EEG response at time *t* at one electrode, or EEG channel *c* can be modeled as

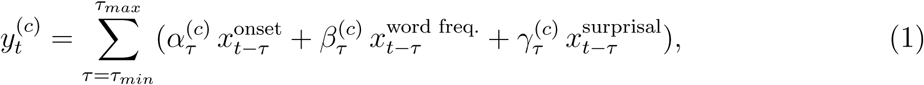

where 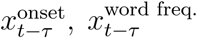, and 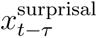 denote the word onset, word frequency, and surprisal feature time series, respectively, at time *t − τ* . 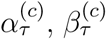, and 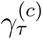 are the corresponding TRF coefficients at time lag *τ* and electrode *c*.

We considered latencies between *τ*_min_ = *−*400 ms and *τ*_max_ = 1.000 ms, capturing both the pre-stimulus phase and the full extent of expected word-evoked cortical responses. Model coefficients were estimated using regularized ridge regression to prevent overfitting, with a regularization parameter *λ* selected from a logarithmically spaced distribution from 10*^−^*^10^ to 10^10^.

Model performance was evaluated using leave-one-trial-out cross-validation. For each left-out trial, the model was fitted on the remaining trials. Prediction accuracy was quantified as the Pearson correlation between the model-predicted and the measured EEG response on the left-out trial, and encoding scores were averaged across cross-validation folds, yielding subject-level estimates.

#### Hierarchical modeling

To assess the unique contribution of each feature over and above lower-level predictors, a hierarchical modeling approach was applied [31, 56]. Features were added cumulatively across three steps: step 1 included word onsets only, step 2 added word frequency, and step 3 further added word surprisal. This ordering reflects a theoretically motivated hierarchy of lexical processing, progressing from purely temporal information (word onset timing), through a context-independent lexical property (word frequency, a proxy for long-term lexical familiarity), to a fully context-dependent measure of predictive processing (surprisal), in line with established accounts of incremental speech comprehension [31]. At each step, the full leave-one-trial-out cross-validation procedure was repeated.

The improvement in prediction accuracy *r* from one step *j* to the next step *j* + 1, Δ*r* = *r_j_*_+1_ *− r_j_*, served as an index of the unique neural variance explained by the added feature, orthogonalized with respect to the preceding predictors [31]. In particular, the gain of word frequency (*r*_2_*−r*_1_) quantified the unique neural encoding of word frequency above and beyond word onsets. The gain of surprisal (*r*_3_ *−r*_2_) quantified the unique neural encoding of surprisal above and beyond word onsets and word frequency.

The final model included all three features simultaneously. This way, we achieved a combined measure of overall neural tracking as well as feature-specific TRF coefficients for each word-level predictor. The associated total gain (*r*_3_ *− r*_1_) captured the combined added benefit of encoding both word frequency and surprisal over the onset-only baseline, and thus served as a broader index of higher-order lexical and predictive processing. All gains were computed separately for the single-speaker and the competing-speaker conditions.

### 2.7 Statistical analysis

#### 2.7.1 Null model

To establish a subject-specific baseline for neural encoding, a null model was computed for each participant alongside the main TRF model. The null model was identical in all respects to the standard model, i.e. it employed the same preprocessing and leave-one-trial-out cross-validation scheme, with the sole exception that all three feature time series (word onset, word frequency, and word surprisal) were time-reversed prior to model fitting. Reversing the temporal order of the features destroys the causal relationship between the stimulus and the EEG response while preserving the overall amplitude distributions and correlation structures of the predictors.

#### 2.7.2 Significance of TRF coefficients

To identify latency windows and electrode clusters in which TRF coefficients significantly exceeded the null model, spatio-temporal cluster-based permutation tests were applied [57]. For each feature (word onset, word frequency, word surprisal) and each listening condition (single speaker, competing speaker attended, competing speaker ignored), the group-level TRF coefficients were compared against the corresponding null-model coefficients. Clusters were formed using a one-tailed F-test threshold (*p <* 0.05), restricted to samples where the TRF coefficients exceeded the null-model coefficients, with spatial adjacency defined by the sensor layout. Cluster-level significance was assessed with 1,000 permutations. To control for the nine simultaneous comparisons (3 features *×* 3 conditions), the cluster-level significance threshold was Bonferroni-corrected to *α* = 0.05*/*9 *≈* 0.0056. Clusters spanning fewer than four electrodes were excluded from further interpretation. For each cluster, we identified peak latencies for topographic visualization. Within each cluster’s temporal window, the peak lag was determined as the sample maximizing the sum of absolute amplitude values across significant channels.

#### 2.7.3 Significance of feature gains

To test whether each individual feature gain was significantly greater than zero, indicating that the corresponding feature led to a reliable above-chance enhancement of the neural encoding, Wilcoxon signed-rank tests were applied to the gain distributions within each cohort (TH, HA, CI) separately. This was done for each of the three gain measures (word frequency gain, surprisal gain, and total gain) and each listening condition (single speaker, competing speaker attended, competing speaker distractor). The *p*-values were corrected for multiple comparisons within each cohort using the Benjamini-Hochberg FDR procedure [58], applied jointly across the three gain measures per cohort and condition.

#### 2.7.4 Selective attention

To examine whether neural encoding scores differentiated the attended from the distractor speaker, we leveraged the competing-speaker condition, in which attended and distractor speakers were presented simultaneously. This allowed a direct within-subject assessment of attentional modulation. For each subject, attended and distractor encoding scores were compared using Wilcoxon signed-rank tests on the paired within-subject differences, as encoding scores were not normally distributed. P-values were FDR-corrected within each cohort across the three model steps. Effect sizes are reported as the rank-biserial correlation (RBC), where values close to 1 indicate that the attended encoding score exceeded the distractor score in nearly all participants. Results are reported as Wilcoxon W-statistic, RBC effect size, and FDR-corrected p-values *p*_fdr_.

### 2.8 Correlation of neural responses to behavior

To examine the relationship between neural encoding gains and behavioral performance, we correlated the three neural gain measures (word frequency gain, surprisal gain, and total gain) with subjective listening effort and speech comprehension scores. For each combination of gain measure and listening condition (single-speaker, competing-speaker attended), we computed one gain value and one corresponding behavioral score (listening effort or comprehension) per participant. Where data from all three cohorts were available, gain and behavioral scores were correlated using repeated-measures correlation [59], as implemented in the pingouin toolbox [60], with cohort (TH, HA, CI) entered as the grouping variable: the model fit a common slope across all participants while allowing a separate intercept per cohort. Comprehension scores were not analyzed for all cohort-by-condition combinations, as performance was at ceiling for TH and HA listeners in both conditions, and for CI users in the single-speaker condition (see Appendix Table 2) [23]. Sufficient variance of comprehension scores was present only for CI users in the competing-speaker condition. For this group, the correlation between neural gain measures and comprehension scores was computed using Pearson correlation, as only a single cohort remained. To control for multiple comparisons, p-values were corrected separately within each of the three resulting families (listening effort single-speaker, listening effort competing-speaker, and comprehension competing-speaker) across the three gain measures, using the Benjamini–Hochberg false discovery rate (FDR) procedure (*n* = 3), as implemented in statsmodels.

### 2.9 Software

All data were analyzed using Python 3.12. Exemplary code is available at https://github.com/Constantin-Jehn/word-level-th-ha-ci.git.

## 3 Results

### 3.1 Neural encoding scores from hierarchical modeling

We assessed the contribution of the linguistic features word onset, word frequency and word surprisal to the neural tracking. Because these linguistic features progress from lower-level (word onset) to higher-level information (word surprisal, which depends on context, we designed a hierarchical model that starts with a baseline model consisting of word onsets only. We then added word frequency in a first step and word surprisal in a second step. Each time we quantified how well the model could explain the neural data through neural encoding scores. To assess whether each additional feature contributed significantly to the predictive power of the model, we computed the gain in encoding score, defined as the difference between encoding scores of successive model steps. We then tested if these gains were significantly above chance level.

#### 3.1.1 Single speaker paradigm

Panel A in Fig. 2 reports the neural encoding scores for the single speaker paradigm. Descriptively, the encoding scores for the different steps of the hierarchical model (left panel) were largest for TH participants, with median values above 0.04, smaller for HA users (median values between 0.03 and 0.04), and lowest for CI users (median values about 0.03). In the first column, baseline encoding scores (word-onset only model) were elevated in TH participants (median = 0.042, IQR = 0.033) compared to HA listeners (median = 0.035, IQR = 0.033), and CI users reached the lowest encoding scores (median = 0.030, IQR = 0.021).

**Figure 2:**
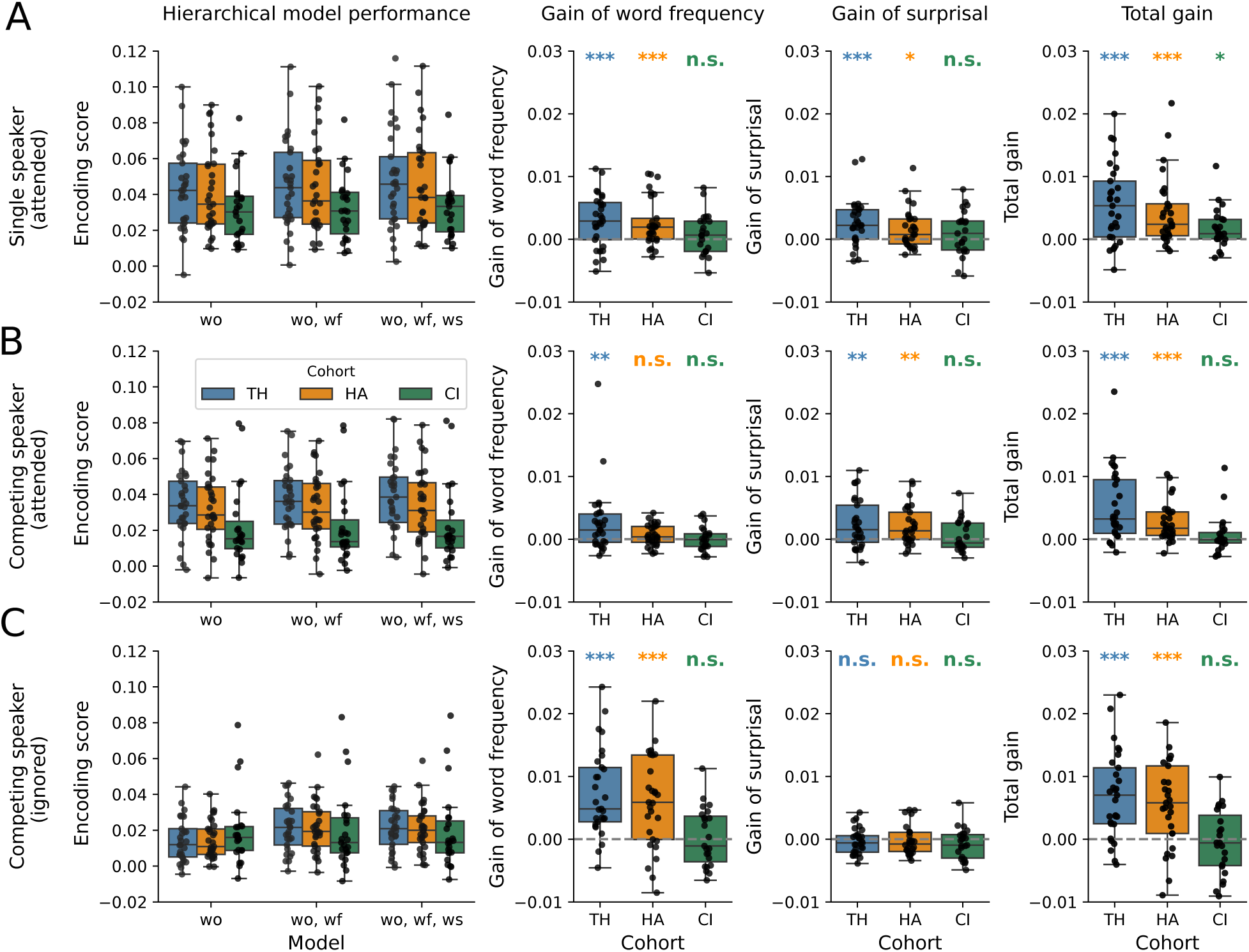
Encoding scores from hierarchical modeling. The left column shows encoding scores for the three hierarchical model stages (wo: word onset only; wo+wf: adding word frequency; wo+wf+ws: adding word surprisal) across the three cohorts. Each data point represents one participant. The remaining columns show the incremental gain achieved by adding word frequency, word surprisal, and their combined effect (wo+wf+ws vs. wo), with stars denoting statistical significance. **A**, In the single-speaker paradigm, TH and HA participants showed overall higher encoding scores than the CI cohort, with significant gains for both word frequency and word surprisal. CI users showed a significant gain only for the full model. **B**, For the attended speaker in the competing-speaker paradigm, word frequency yielded a significant gain only in the TH group, while word surprisal produced significant gains in both TH and HA participants. CI users showed no significant gain for either feature. **C**, Encoding scores for the distractor stream were overall reduced in TH and HA participants across all model stages; nevertheless, both groups showed significant gains when word frequency was added. As expected, no group exhibited a significant surprisal gain for the distractor speaker (***: *p <* 0.001, **: *p <* 0.01, *: *p <* 0.05, n.s.: *p ≥* 0.05).

We now computed the gains, i.e., the differences in encoding scores, between two successive steps of hierarchical modelling. Adding word frequency to the baseline model of word-onset only yielded a significant encoding gain at the group level in both the TH (*W* = 66, *RBC* = 0.70, *p*_FDR_ = 0.001) and HA (*W* = 66, *RBC* = 0.70, *p*_FDR_ = 0.001) cohorts, whereas no significant gain was observed in CI users (*W* = 105, *RBC* = 0.24, *p*_FDR_ = 0.329; Panel A - Gain of word frequency).

Further adding word surprisal to the model produced an additional significant gain in both TH (*W* = 65, *RBC* = 0.70, *p*_FDR_ = 0.001) and HA (*W* = 105, *RBC* = 0.52, *p*_FDR_ = 0.018) participants, but not in CI users (*W* = 99, *RBC* = 0.28, *p*_FDR_ = 0.278; Panel A - Gain of surprisal). Consequently, the total gain of the full model over the baseline model (Panel A - Total gain) was highly significant for both TH (*W* = 40, *RBC* = 0.82, *p*_FDR_ *<* 0.001) and HA (*W* = 32, *RBC* = 0.85, *p*_FDR_ *<* 0.001) participants. Notably, this gain also reached significance for the CI users (*W* = 51, *RBC* = 0.63, *p*_FDR_ = 0.010), indicating that word-level neural tracking is present even in cochlear implant users, albeit with a considerably smaller effect size than observed in TH and HA groups.

#### 3.1.2 Competing speaker paradigm: attended speaker

Panel B in Fig. 2 depicts the encoding scores for the attended speaker in the competing speaker paradigm. Descriptively, baseline encoding scores were again elevated in TH (median = 0.034, IQR = 0.024) and HA (median = 0.029, IQR = 0.024) participants relative to CI users (median = 0.015, IQR = 0.015), consistent with the single speaker paradigm (left panel).

Adding word frequency to the model (Panel B - Gain of word frequency) yielded a significant gain only in the TH group (*W* = 128, *RBC* = 0.63, *p*_FDR_ = 0.005), while HA (*W* = 81, *RBC* = 0.37, *p*_FDR_ = 0.126) and CI users (*W* = 137, *RBC* = *−*0.07, *p*_FDR_ = 0.874) showed no significant benefit at this step. However, adding surprisal (Panel B - Gain of surprisal) produced a significant gain in both TH (*W* = 111, *RBC* = 0.59, *p*_FDR_ = 0.008) and HA (*W* = 89, *RBC* = 0.67, *p*_FDR_ = 0.003) groups, indicating that a robust context-related processing to the attended speaker was preserved under competing speech conditions in both typical-hearing and hearing-aid users. CI users again showed no significant surprisal gain (*W* = 71, *RBC* = 0.20, *p*_FDR_ = 0.550). The total gain over the word-onset baseline (right panel) was highly significant for TH (*W* = 135, *RBC* = 0.87, *p*_FDR_ *<* 0.001) and HA (*W* = 29, *RBC* = 0.89, *p*_FDR_ *<* 0.001) groups (Panel B - Total gain). In contrast, CI users showed no significant total gain (*W* = 24, *RBC* = *−*0.02, *p*_FDR_ = 0.941).

#### 3.1.3 Competing speaker paradigm: distractor speaker

Panel C in Fig. 2 shows the encoding scores for the ignored speaker in the competing speaker paradigm. Descriptively, compared to the other two previous conditions, baseline word-onset encoding scores were markedly reduced in both TH (median = 0.012, IQR = 0.016) and HA (median = 0.011, IQR = 0.014) participants, approaching the levels observed in CI users (median = 0.016, IQR = 0.013; left panel). The ignored speaker thus elicits a substantially weaker word-onset response compared to the attended speaker for the TH and HA group, but not for the CI group. Adding word frequency to the model (Panel C - Gain of word frequency) still yielded a highly significant gain in both TH (*W* = 131, RBC = 0.91, *p*_FDR_ *<* 0.001) and HA (*W* = 19, RBC = 0.72, *p*_FDR_ = 0.001) participants, while CI users showed no significant gain (*W* = 61, RBC = 0.05, *p*_FDR_ = 0.846). Despite the large incremental gain of adding word frequency, the absolute encoding scores after this step remained markedly lower in the distractor condition than in the attended scenario for both TH (median = 0.022, IQR = 0.020 vs. median = 0.036, IQR = 0.024) and HA (median = 0.019, IQR = 0.019 vs. median = 0.030, IQR = 0.025), underscoring the substantial effect of selective attention on word frequency tracking.

In contrast, no group showed a significant gain when surprisal was added (Panel C - Gain of surprisal; TH: *W* = 89, RBC = *−*0.24, *p*_FDR_ = 0.341; HA: *W* = 165, RBC = *−*0.25, *p*_FDR_ = 0.341; CI: *W* = 163, RBC = *−*0.36, *p*_FDR_ = 0.256). As a consequence of the robust word frequency contribution, the total gain remained highly significant for TH (*W* = 125, RBC = 0.86, *p*_FDR_ *<* 0.001) and HA (*W* = 30, RBC = 0.75, *p*_FDR_ = 0.001) groups, whereas CI users showed no significant total gain (*W* = 55, RBC = *−*0.09, *p*_FDR_ = 0.798; right panel).

#### 3.1.4 Selective attention

We further analyzed whether the encoding scores of the hierarchical models were affected by selective attention in the competing talker condition (Fig. 2B, C). To this end, for each subject, we compared the encoding scores obtained for the attended talker to those for the ignored talker using Wilcoxon signed-rank tests on the paired within-subject differences, for all models within the three cohorts.

##### TH individuals

TH individuals showed a highly significant difference between attended and ignored encoding scores at every model step. At the word onset level, attended encoding scores were markedly higher than ignored scores (attended: median = 0.034, IQR = 0.024; ignored: median = 0.012, IQR = 0.016; *W* = 13, RBC = 0.940, *p*_fdr_ *<* 0.001). This difference persisted after adding word frequency (attended: median = 0.036, IQR = 0.024; ignored: median = 0.022, IQR = 0.020; *W* = 9, RBC = 0.959, *p*_fdr_ *<* 0.001), and word surprisal (attended: median = 0.039, IQR = 0.025; ignored: median = 0.021, IQR = 0.019; *W* = 2, RBC = 0.991, *p*_fdr_ *<* 0.001). Notably, while the absolute difference between attended and ignored scores decreased slightly when adding model steps, the RBC increased monotonically, indicating that attentional modulation became more consistent across participants as more linguistic complexity was included in the model.

##### HA users

Similar to TH individuals, HA listeners showed a highly significant difference between attended and ignored encoding scores at every model step. At the word onset level, attended encoding scores were substantially higher than ignored scores (attended: median = 0.029, IQR = 0.024; ignored: median = 0.011, IQR = 0.014; *W* = 28, RBC = 0.871, *p*_fdr_ *<* 0.001). This difference persisted after adding word frequency (attended: median = 0.030, IQR = 0.025; ignored: median = 0.019, IQR = 0.019; *W* = 29, RBC = 0.867, *p*_fdr_ *<* 0.001) and word surprisal (attended: median = 0.031, IQR = 0.027; ignored: median = 0.020, IQR = 0.015; *W* = 25, RBC = 0.885, *p*_fdr_ *<* 0.001). In contrast to TH individuals, the RBC remained stable when adding model steps rather than increasing monotonically, suggesting that while attentional modulation was robust and consistent throughout, the progressive addition of linguistic features did not further sharpen the attended–ignored distinction in HA listeners.

##### CI users

CI users showed a markedly different pattern from both TH and HA listeners. At the word onset level, attended and ignored encoding scores were virtually identical (attended: median = 0.015, IQR = 0.015; ignored: median = 0.016, IQR = 0.013; *W* = 106, RBC = 0.232, *p*_uncorr_ = 0.345, *p*_fdr_ = 0.345), indicating no reliable attentional modulation of the word onset response. Similarly, adding word frequency did not yield a significant attended–ignored difference (attended: median = 0.014, IQR = 0.015; ignored: median = 0.013, IQR = 0.019; *W* = 105, RBC = 0.239, *p*_uncorr_ = 0.329, *p*_fdr_ = 0.345). For the full model, including word surprisal, a nominally significant difference, with medium effect size, emerged between attended and ignored scores (attended: median = 0.017, IQR = 0.015; ignored: median = 0.013, IQR = 0.018; *W* = 72, RBC = 0.478, *p*_uncorr_ = 0.045). However, this difference did not survive FDR correction (*p*_fdr_ = 0.135). Overall, CI users thus showed no significant attentional modulation of the neural encoding scores at the word-level at any model step.

### 3.2 Temporal Response Function

We next examined the Temporal Response Functions (TRFs) of the full model, incorporating word onset, word frequency, and word surprisal. While encoding scores provide a global measure of model performance, TRFs reveal the timing of neural responses and, through their topographies, give indication of the underlying neural generators. Importantly, encoding scores that are nonsignificant at the group level do not preclude significant responses at specific time lags or individual channels.

Figs. 3, 4, and 5 show the TRFs for the TH participants, the HA users, and the CI users, respectively. Channels and time lags at which the TRFs significantly exceed the null distribution (spatio-temporal cluster permutation test) are rendered in bold. Topographies adjacent to the butterfly plots show the scalp distribution at peak latencies, and only significant channels are shown as dots in the topographies.

**Figure 3:**
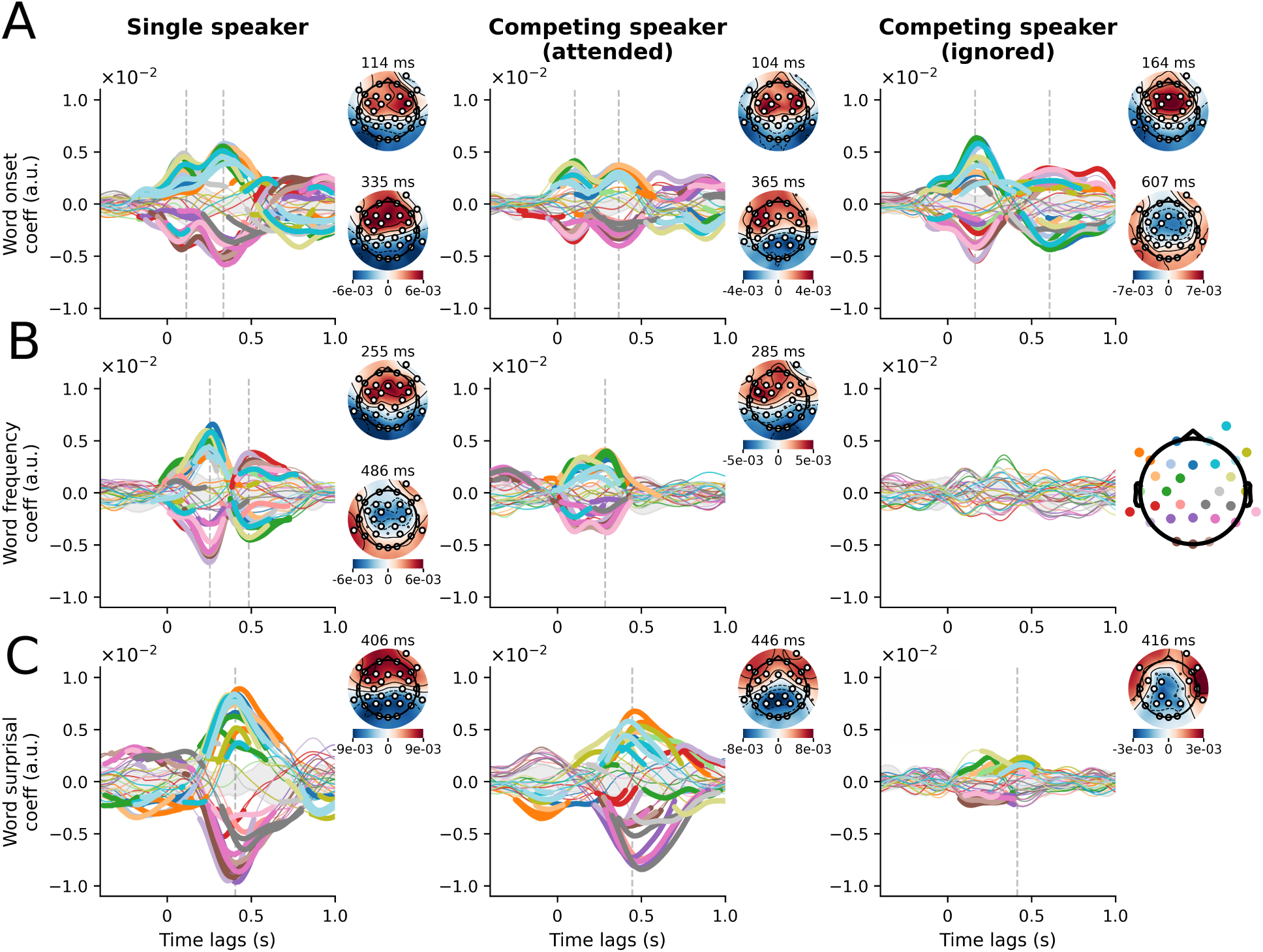
Temporal response functions (TRFs) of the full model for typical-hearing (TH) participants. The three columns correspond to the three listening conditions: single speaker, competing speaker (attended), and competing speaker (ignored). Scalp topographies are shown for significant amplitude peaks. **A**, Word onset responses, peaking at 114 and 335 ms with fronto-central topographies. Responses to the attended stream in the competing speaker condition follow a similar pattern at reduced amplitude. Responses to the ignored speaker are delayed. **B**, Word frequency responses first peak at 254 ms, later than the first word onset peak. Responses are reduced in the competing speaker condition and largely absent for the distractor stream. **C**, Neural responses to word surprisal peak at 405 ms with a dipolar scalp topography, consistent with the N400 response. The response is preserved in the competing speaker condition for the attended talker, whereas the ignored speech stream yielded a weaker response with fewer significant channels. Bold channels and latencies indicate significant spatio-temporal clusters; topographies are shown at peak latencies.

**Figure 4:**
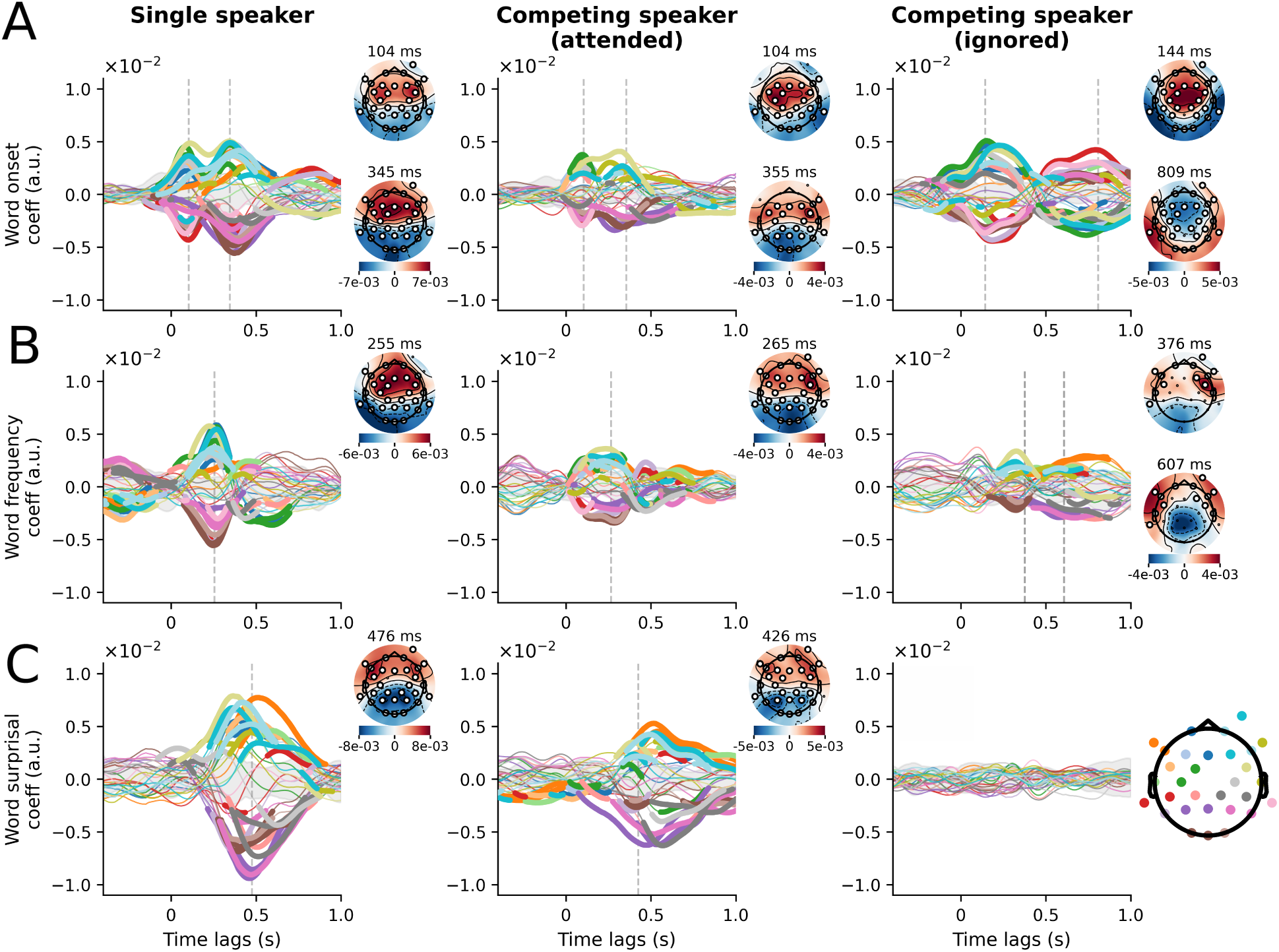
Temporal response functions (TRFs) of the full model for hearing aid (HA) users. The three columns correspond to the three listening conditions: single speaker, competing speaker (attended), and competing speaker (ignored). **A**, Word onset responses peak at 104 and 345 ms with fronto-central topographies, closely matching the TH group. Both peaks are preserved in the competing speaker condition at reduced amplitude. **B**, Word frequency responses peak at 255 ms, attenuated but preserved in latency for the attended stream in the competing speaker condition. The distractor stream yielded fewer significant channels. **C,** Word surprisal responses peak at 476 ms with a dipolar topography, consistent with the N400. A clear response was preserved for the attended stream, whereas the distractor stream produced no significant response. Bold channels and latencies indicate significant spatio-temporal clusters. Topographies are shown at peak latencies.

**Figure 5:**
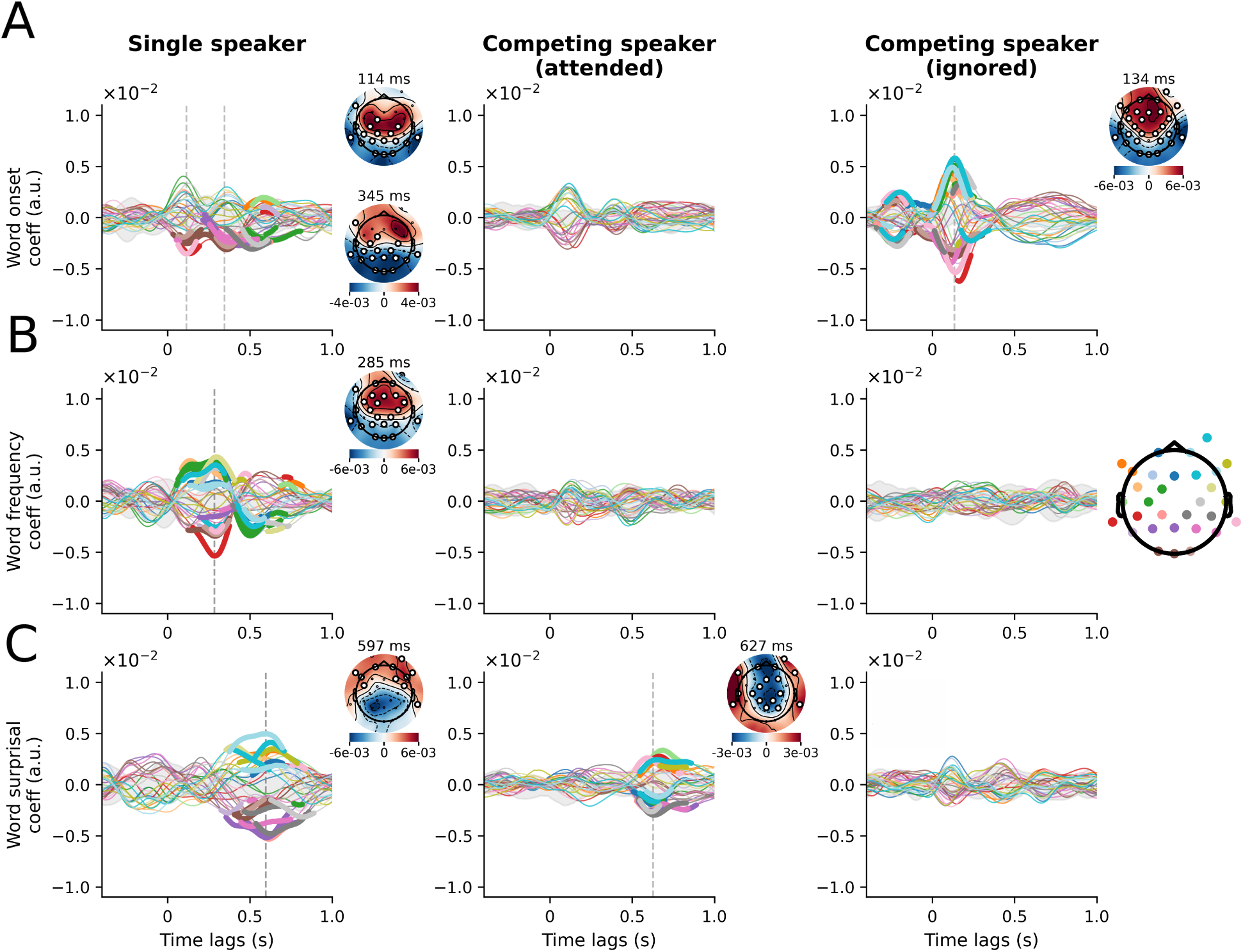
Temporal response functions (TRFs) of the full model for cochlear implant (CI) users. The three columns correspond to the three listening conditions: single speaker, competing speaker (attended), and competing speaker (ignored). **A**, Word onset responses are weaker than in both other cohorts, with fewer significant channels in the single speaker condition and no significant response in the competing speaker condition. **B**, Word frequency responses peak at 284 ms in the single speaker condition, comparable in latency to the TH and HA groups. They are absent in the competing speaker condition for both speech streams. **C**, Word surprisal responses peak at 597 ms, later and less pronounced than in both other cohorts. A significant response persists for the attended stream at 627 ms, whereas the ignored stream produced no significant cluster. Bold channels and latencies indicate significant spatio-temporal clusters. Topographies are shown at peak latencies.

#### 3.2.1 Typical-hearing participants

Fig. 3 A shows the word onset TRFs across the three listening conditions. In the single speaker condition, two significant peaks emerged at 114 and 335 ms (mean amplitude 0.004 a.u. and 0.005 a.u., respectively). across 23 significant channels, both with frontocentral topographies. In the competing speaker condition, the attended stream produced a similar pattern at delays of 104 and 365 ms, with reduced amplitude across 18 and 22 significant channels (mean amplitude both 0.003 a.u., respectively). The ignored stream produced a stronger first peak at a delay of 164 ms across 25 significant channels (mean amplitude = 0.004 a.u.) and a late response at 607 ms across 20 channels (mean amplitude = 0.003 a.u.), both with fronto-central topographies.

The TRFs for word frequency are depicted in Fig. 3 B. In the single speaker condition, two significant peaks emerged at delays of 255 and 486 ms across 21 and 17 significant channels (mean amplitude = 0.004 and 0.003 a.u., respectively). The first peak at 255 ms was later than the word onset peak at 114 ms, indicating processing at a higher level of the auditory hierarchy. The topography was fronto-central. In the competing speaker condition, the attended stream produced a significant peak at 285 ms across 20 channels (mean amplitude = 0.003 a.u.), attenuated relative to the single speaker condition. The ignored stream yielded no significant clusters.

Fig. 3C shows the TRFs for word surprisal. In the single speaker condition, a significant peak emerged at a latency of 406 ms across 27 channels (mean amplitude = 0.007 a.u.), later than both the word onset and word frequency peaks. This latency is consistent with the N400 component known from ERP research. The topography showed a dipolar structure with significant channels across the entire scalp. In the competing speaker condition, the attended stream produced a peak at 446 ms across 23 channels (mean amplitude = 0.005 a.u.) with a similar topography. The distractor stream yielded only five significant channels at a delay of 416 ms (mean amplitude = 0.003 a.u.), indicating a strong effect of attention on the surprisal response.

#### 3.2.2 Hearing aid users

Fig. 4 shows the TRFs for hearing aid (HA) users. Word onset responses (panel A) closely resembled those of TH participants. In the single speaker condition, two significant frontocentral peaks emerged at delays of 104 and 345 ms across 22 and 24 significant channels (mean amplitude = 0.003 and 0.005 a.u., respectively). Both peaks were preserved in the attended stream of the competing speaker condition at latencies of 104 and 355 ms, with reduced amplitude across 17 and 10 significant channels (mean amplitude = 0.003 and 0.003 a.u., respectively). The distractor stream produced responses at 144 and 809 ms across 21 and 16 significant channels (mean amplitude = 0.003 and 0.003 a.u.), consistent with the TH group.

Word frequency responses (panel B) showed a single peak at 255 ms across 21 significant channels (mean amplitude = 0.004 a.u.) in the single speaker condition. In the competing speaker scenario, this peak was attenuated but preserved in latency at 265 ms across 15 channels (mean amplitude = 0.003 a.u.) for the attended stream. The ignored stream yielded only five significant channels at 376 ms and seven channels at 607 ms, indicating a markedly weaker response.

Word surprisal responses (panel C) peaked at 476 ms across 23 significant channels (mean amplitude = 0.005 a.u.) in the single speaker condition, consistent with an N400 response. In the competing speaker condition, a significant peak at 426 ms was preserved for the attended stream across 15 channels (mean amplitude = 0.003 a.u.), though with reduced amplitude. The distractor stream produced no significant response, indicating that no suprisal-related processing was detected for the unattended speaker.

#### 3.2.3 Cochlear Implant users

Fig. 5 shows the TRFs for cochlear implant (CI) users. Word onset responses (panel A) were weaker than in both other cohorts already in the single-speaker condition, with significant peaks at delays of 114 ms and 345 ms, spanning five and nine channels, respectively (both mean amplitude = 0.003 a.u.). In the competing speaker condition, the attended stream showed a peak descriptively, which did not survive cluster forming and statistical testing. The distractor stream, however, yielded a significant peak at a latency of 134 ms across 16 channels (mean amplitude = 0.005 a.u.).

Word frequency responses (panel B) showed a single significant peak at 285 ms across nineteen significant channels (mean amplitude = 0.004 a.u.) in the single speaker condition, comparable in latency to the TH and HA groups. In the competing speaker condition, neither the attended nor the distractor stream produced a significant response.

Word surprisal responses (panel C) showed a significant peak at 597 ms across nine channels (mean amplitude = 0.004 a.u.) in the single speaker condition. This latency is substantially later than the delays of 406 and 476 ms of the peaks observed in TH and HA participants, respectively. In the competing speaker condition, a significant response persisted for the attended stream at 627 ms across 15 channels (mean amplitude = 0.003 a.u.), whereas the distractor stream produced no significant cluster. The presence of a significant attended response alongside a nonsignificant distractor response evidences a weak but detectable attentional modulation of context-dependent word processing in CI users.

### 3.3 No significant relation to behavior

We further analyzed the potential relation of the neural tracking of the linguistic features to behavioral outcomes. Specifically, we correlated word frequency gain, surprisal gain, and the total gain reported in Fig. 2 to the subjective listening effort on the ACALES scale, and to the comprehension scores in the competing speaker paradigm for the CI users. TH participants and HA users were excluded from the latter analysis as their speech comprehension was at ceiling (i.e., they answered almost all questions correctly) and thus did not have sufficient variance for a useful analysis.

Repeated-measures correlation revealed no significant relationship between any of the neural gain measures and subjective listening effort in the single-speaker condition (Fig. 6A). All correlation coefficients were near zero and the associated *p*-values were well above 0.05 (*r* = 0.03 for word frequency gain, *r* = *−*0.08 for surprisal gain, *r* = *−*0.03 for total gain; *p*_fdr_ = 0.794 for all three measures).

**Figure 6:**
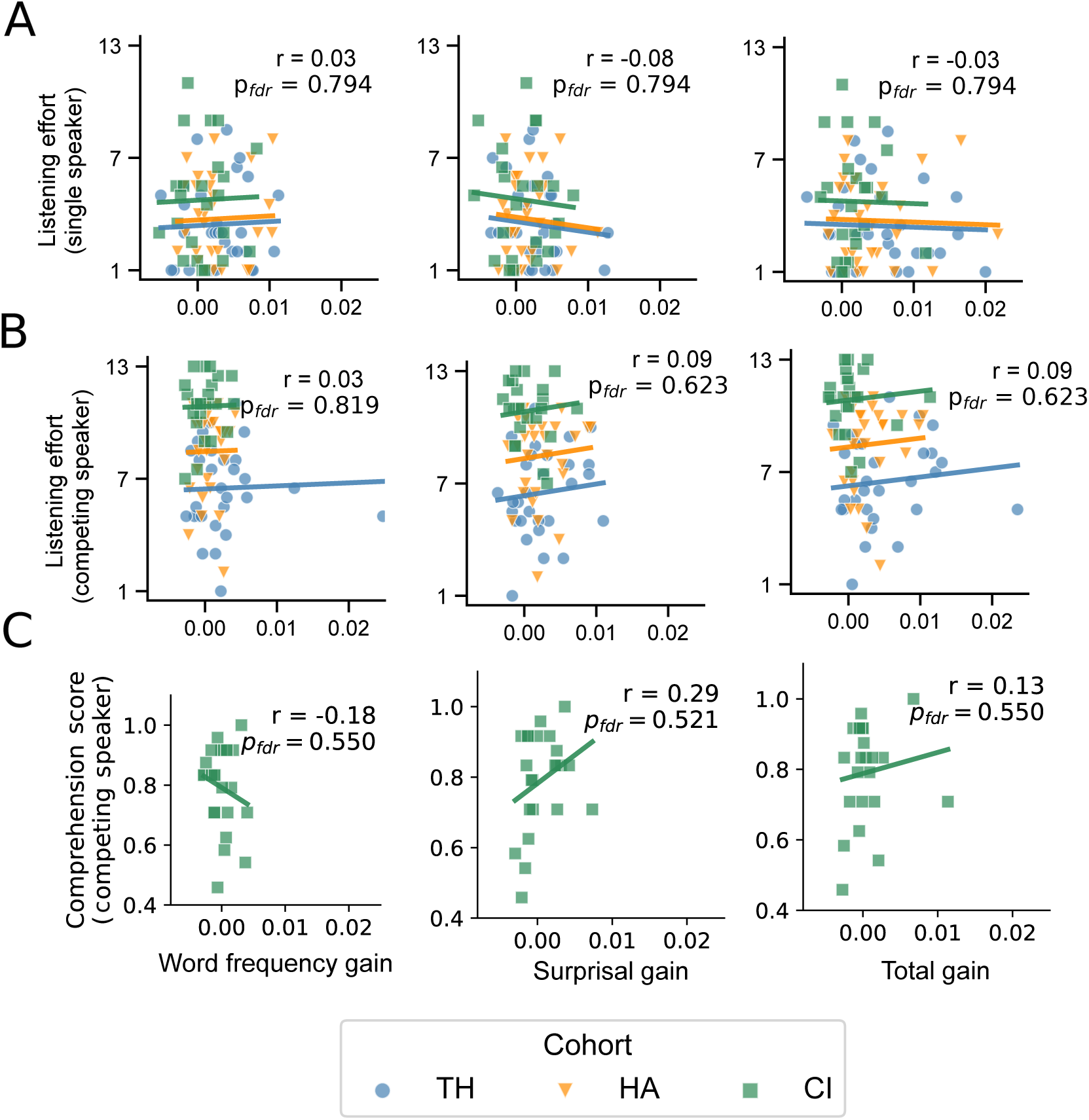
Correlation of neural gains with behavior. Listening effort was analyzed using repeated-measures correlation, with cohort entered as the grouping variable to control for systematic between-cohort differences while estimating the common within-cohort association. Comprehension scores showed sufficient variance only in the CI cohort; Pearson correlation was therefore used for this analysis. **A**, None of the neural gain measures significantly predicted perceived listening effort in the single-speaker condition. **B**, Similarly, no significant relationship was found between any neural gain measure and subjective listening effort in the competing-speaker condition. **C**, In the CI cohort, no significant correlation was observed between any neural gain measure and speech comprehension scores in the competing-speaker condition. Note the restricted range of neural gain values across participants.

Consistent with the single-speaker condition, repeated-measures correlation revealed no significant relationship between any of the neural gain measures and subjective listening effort in the competing-speaker condition either (Figure 6B). Correlation coefficients remained near zero with large associated *p*-values (word frequency gain: *r* = 0.03; *p*_fdr_ = 0.819, surprisal gain: *r* = 0.09; *p*_fdr_ = 0.623, total gain: *r* = 0.09; *p*_fdr_ = 0.623).

Pearson correlation within the CI group revealed no significant relationship between any of the neural gain measures and speech comprehension scores in the competing-speaker condition (Figure 6C). Although correlation coefficients were somewhat larger in absolute value than those observed for listening effort, the *p*-values were far too high for statistical significance (word frequency gain: *r* = *−*0.18; *p*_fdr_ = 0.550, surprisal gain *r* = 0.29 *p*_fdr_ = 0.521, total gain *r* = 0.13; *p*_fdr_ = 0.550).

## 4 Discussion

In the present study, we investigated neural tracking of word-level linguistic information in hearing aid and cochlear implant users, alongside an age-matched, typical-hearing control group. This constitutes the first comparative investigation of word-level tracking across these participant groups, and the first study of word-level tracking in CI users.

### Comparable neural tracking in HA users and TH participants

Overall, the HA group exhibited neural tracking of linguistic features that closely resembled that of the TH group, although for competing speech, we observed some feature-specific differences between these two groups (Figure 2). In both groups, word frequency and surprisal each contributed a significant gain in encoding score during single-speaker listening. Also, for both groups, the combined model yielded a significant total gain in both single and competing listening conditions.

We further found a similar effect of selective attention on the encoding scores in both participant groups, as reflected in the difference between attended and ignored encoding scores. For both TH and HA listeners, this difference was highly significant at every hierarchical model step, with comparably large effect sizes.

A different pattern emerged between the groups for the encoding gain attributable to word frequency (i.e., the increase in encoding score from adding word frequency as a hierarchical model step) in the competing-speaker condition, where this gain reached significance only in the TH group, and not in the HA group.

These findings suggest that HA users do not exhibit a general impairment of word-level linguistic tracking. This overall preservation during single-speaker listening differs from the findings of Bolt and Giroud [33], who reported decreased encoding accuracy for a word-level segmentation model based on word onsets with increasing hearing impairment in older adults. However, their sample did not include hearing aid users and consisted predominantly of older adults with normal hearing or unaided mild hearing impairment, limiting direct comparison with the present HA group. In the present study, the only apparent deviation from the TH pattern concerned word frequency tracking (i.e., the gain in encoding score in the hierarchical model) under competing speech for the attended speaker, raising the possibility that this aspect of lexical processing may be particularly susceptible to the additional demands imposed by a competing speaker.

### Significant surprisal response in CI users

Our analysis of TRF waveforms for word surprisal revealed significant responses in CI users for the single-talker condition and for the attended stream in the competing-talker condition (Figure 5). This constitutes the first evidence of significant neural tracking of word surprisal in CI users. However, the responses were less widespread and descriptively smaller than in the TH and HA groups. Moreover, responses peaked at later latencies (597 ms and 627 ms) than in the other cohorts.

Comparing these results to the ERP study by Hahne et al. [61], the delayed TRF surprisal responses parallel previous ERP findings of prolonged N400 latencies during semantic sentence processing in adult CI users [61, 62]. These findings provide converging evidence for delayed context-dependent semantic integration in this population. The present study extends this evidence from experimentally constructed sentences containing semantic violations to natural continuous speech, showing that delayed sensitivity to the contextual predictability of individual words can also be detected during the comprehension of an unfolding narrative.

### Encoding gain of the linguistic model in CI users

The analysis of encoding scores from the hierarchical model in the single-talker paradigm revealed no significant improvement in encoding score for the CI group when adding word frequency or word surprisal individually. However, the full model integrating all three features (word onset, word frequency, and word surprisal) reached a significantly higher encoding score than the baseline model. This suggests that lexical and contextual information jointly contribute to the neural responses of CI users, although the unique contribution of neither feature reached significance on its own.

We also found no improvement at the level of individual features in the competing-speaker paradigm (attended stream): neither the addition of word frequency nor of word surprisal yielded a significant gain in encoding score for CI users. Unlike in the single-speaker condition, we also found no significant improvement for the combined model. This suggests that the presence of a competing speaker reduces the robustness of neural tracking of lexical and contextual level information in CI users.

This finding stands in seeming contrast to the significant surprisal TRF response we observed for the attended stream in the competing-speaker condition, which provides evidence that context-dependent word processing remains detectable while competing speech is present. We argue, however, that these two observations are not contradictory, but reflect a documented distinction between encoding-score-based and TRF-based measures of neural tracking. The encoding score and the gain measures derived from it summarize prediction accuracy across the entire response window and across all scalp channels. A significant spatio-temporally localized response in the word surprisal TRF does not necessarily translate into a detectable improvement in the aggregate metric. A similar dissociation was reported by Mesik et al. [63], who found that attentional modulation was more pronounced in their TRFs than in their model-fit contributions.

### Reduced attentional modulation in CI users

Analyzing the difference in encoding scores between attended and ignored voices in the competing speaker condition showed a pronounced attention effect in the TH and HA groups across the three steps of hierarchical modeling. In contrast, the CI group showed no significant attentional modulation for the baseline model (word onset) nor for the second step (word onset and word frequency). When word surprisal was added to the model in the third step, a medium effect size was observed for the combined model. Yet, the effect was not significant after FDR correction.

Consistent with attenuated responses to the speech envelope in cochlear implant users [23, 24], the CI group thus showed impaired selectivity between attended and distractor speech to word-level features. Nonetheless, the significant peak in the TRFs for surprisal for the attended talker, but not for the ignored voice, suggests that surprisal is tracked selectively in this group, supporting speech understanding in competing-speaker scenarios. The markedly reduced magnitude of this effect, however, mirrors the well-documented real-life difficulties CI users face in understanding speech in noisy, multi-talker environments [10]. Those difficulties were also reflected in decreased comprehension scores of CI users compared to TH and HA listeners in the analyzed dataset.

### Strong word onset responses in distractor TRFs

Surprisingly, early word onset responses at latencies of 120–180 ms were larger for the distractor speaker than for the attended speaker across all three cohorts (Figures 3, 4, 5). In the TH group, this response to the ignored talker involved a larger number of significant channels (25 vs. 18) and a larger mean amplitude (0.004 vs. 0.003 a.u.) than the attended response. The HA group showed a similar pattern, with more significant channels (21 vs. 17) and a modestly larger amplitude (0.032 vs. 0.026 a.u.) for the distractor. In the CI group, this effect was most pronounced: the early word onset response reached significance only for the distractor speaker (16 channels; mean amplitude = 0.021 a.u.), with no corresponding significant response to the attended speaker.

A comparable dominance of onset encoding for the ignored over the target speaker was recently reported in hearing-impaired listeners by Carta et al. [34]. One of their interpretations was that the robust representation of distractor phoneme onsets is a neural indicator for the higher distractibility reported in this population. Our data suggest that this phenomenon is not specific to hearing impairment, as the same distractor-dominant onset response is present in typical-hearing listeners. Our data more closely support the alternative interpretation proposed by Carta et al. [34]: preserving a strong structural representation of the distractor speaker enables the listener to switch the focus of attention, as needed.

### Absence of neuro-behavior correlations

Our correlation analysis revealed no significant relationship between the neural gains in encoding scores for word frequency, surprisal, and their combined total, and the behavioral measures of listening effort and comprehension. We note three limitations that likely constrained this analysis. First, comprehension scores were at ceiling for the TH and HA cohorts across all conditions, leaving little room for these scores to covary with neural measures, so they were not analyzed. Second, the CI group showed only moderate gains from word-level tracking, restricting the range of variance available on the neural side of the analysis. Third, since the comprehension questions assessed memory for the overall content of each two-minute trial as a whole, they offered only a coarse measure of comprehension rather than a fine-grained index sensitive to word-by-word processing.

In a paradigm similar to ours, Mesik et al. [63] found a non-significant positive trend between speech comprehension and the fit of a word-level model in parietal channels for typical-hearing individuals. Using a more fine-grained measure of speech comprehension, Barchet et al. [64] found that enhanced neural tracking of word frequency and word surprisal for the target speaker significantly predicted higher speech comprehension: In their study, comprehension was measured as the proportion of words correctly repeated from two sentences, offering a more fine-grained, word-level measure of understanding than the triallevel comprehension questions used here. With 240 trials per participant, their study also afforded substantially greater statistical power for neuro-behavioral analyses than the 20 trials available in our data.

## 5 Limitations

As this analysis draws on the same dataset as our previous study [23], several limitations discussed there apply here as well. These include the CI cohort’s inclusion criterion of scoring at least 60% on the Freiburg monosyllabic word recognition test in quiet: while this represents a common performance level among CI users [65, 66], poorer performers are underrepresented in the dataset. However, this was intentional as the competing talker paradigm is already an exhausting experience for well-performing CI users. Data on binaural performance in CI users are also missing, which would clarify to what extent these listeners benefit from spatial release from masking.

Analyzing EEG data from CI users additionally requires removing electrical stimulation artifacts. However, we note that responses to linguistic features occur at latencies *≥* 100 ms and are therefore less susceptible to these artifacts. The surprisal response, for instance, peaked at a relatively long delay of 600 ms, making it unlikely to be meaningfully affected by stimulation artifacts. Finally, the use of participants’ personal hearing aids and cochlear implants increased the ecological validity of our findings, but also introduced variability across participants due to differences in device type and individual fitting parameters; in CI users, different stimulation strategies may have introduced additional variability. We argue, however, that using the optimal fitting parameters for each participant ensures that they are able to perform to the best of their abilities.

## 6 Conclusion

This work presented the first comparative study of neural tracking of word-level features across CI users, HA users, and an age-matched TH control group. Neural tracking was observed both for a single-speaker as well as a competing speaker condition presented in free-field with hearing-impaired individuals using their own hearing devices and settings. The neural tracking of linguistic features was very similar in the HA and the TH participants. In contrast, CI users exhibited much weaker and delayed neural responses. However, our study notably provides the first evidence of significant neural tracking of word surprisal in CI users in response to continuous speech.

The competing speech paradigm allowed us to assess the effect of selective attention to one of two competing narrators. HA and TH participants showed large attention modulation of their neural tracking to all three linguistic features words onset, word frequency, and word surprisal. CI users only exhibited an attentional effect regarding word surprisal: TRFs for the attended talker showed significant neural tracking of word surprisal, while TRFs for the ignored talker did not. These results suggest that neural responses to word surprisal are most informative about selective attention, including in CI users.

Together, these findings refine our understanding of how lexical and content-dependent speech information is neurally processed in hearing aid and cochlear implant users under single and competing speaker conditions.

## Author contributions

Constantin Jehn and Alina Schüller performed the formal analysis, visualized the results, and wrote the manuscript. Niki Katerina Vavatzanidis, Anja Hahne, and Tobias Reichenbach supervised the research, interpreted the results, and edited the manuscript.

## Declaration of generative AI and AI-assisted technologies in the writing process

During the preparation of this work, the authors used Claude in order to improve readability and language. After using this tool, the authors reviewed and edited the content as needed and take full responsibility for the content of the published article.

## Data availability

The data used for this study are publicly available. We used three publicly available datasets:

The data of the cochlear implant group are available under https://doi.org/10.5281/zenodo.17952844.

The data of the hearing-aided group can be obtained from https://doi.org/10.5281/zenodo.17927767.

The data of the typical-hearing group are available from https://doi.org/10.5281/zenodo.17952231.

## A Appendix: Behavioral data

**Table 2:** Behavioral performance by group. Data are published in [23].

| Group | Listening Effort<br>(Single Speaker) | Listening Effort<br>(Competing Speaker) | Comprehension<br>(Single Speaker) | Comprehension<br>(Competing Speaker) |
|---|---|---|---|---|
| TH (*n* = 29) | 3.45 ± 2.23 | 6.50 ± 2.33 | 0.983 ± 0.037 | 0.990 ± 0.029 |
| HA (*n* = 29) | 3.74 ± 2.27 | 8.47 ± 2.34 | 0.987 ± 0.031 | 0.977 ± 0.043 |
| CI (*n* = 24) | 4.79 ± 2.86 | 10.83 ± 1.61 | 0.979 ± 0.040 | 0.798 ± 0.135 |

Values are mean ± SD. Listening effort was assessed using the ACALES scale (1–13). Comprehension scores are proportion correct (0–1).

